## Supplementary figures and images for "Dietary sulfur amino acid restriction elicits a cold-like transcriptional response in inguinal but not epididymal white adipose tissue of male mice"

### Supplemental figure 1

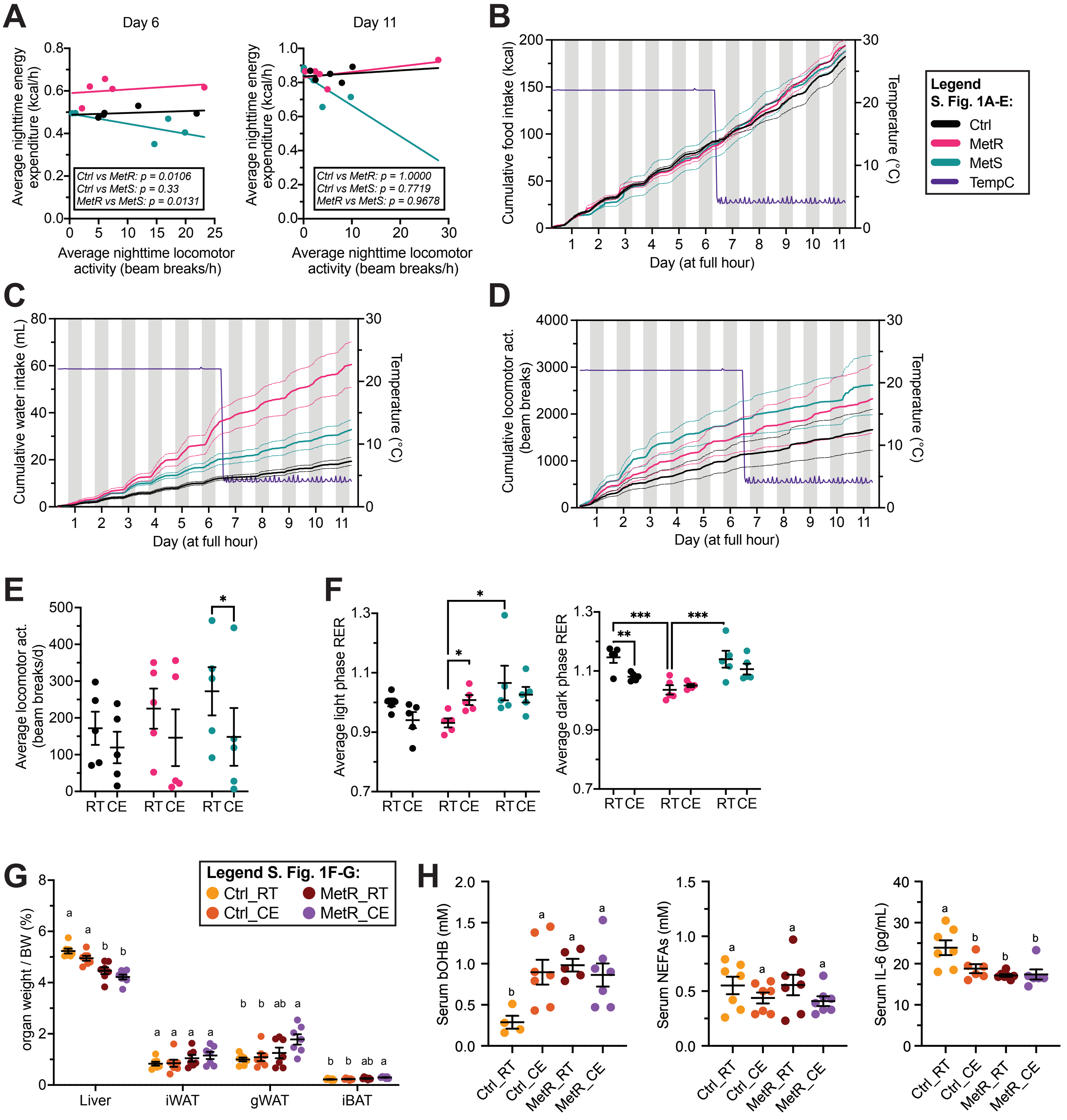

### Supplemental figure 2

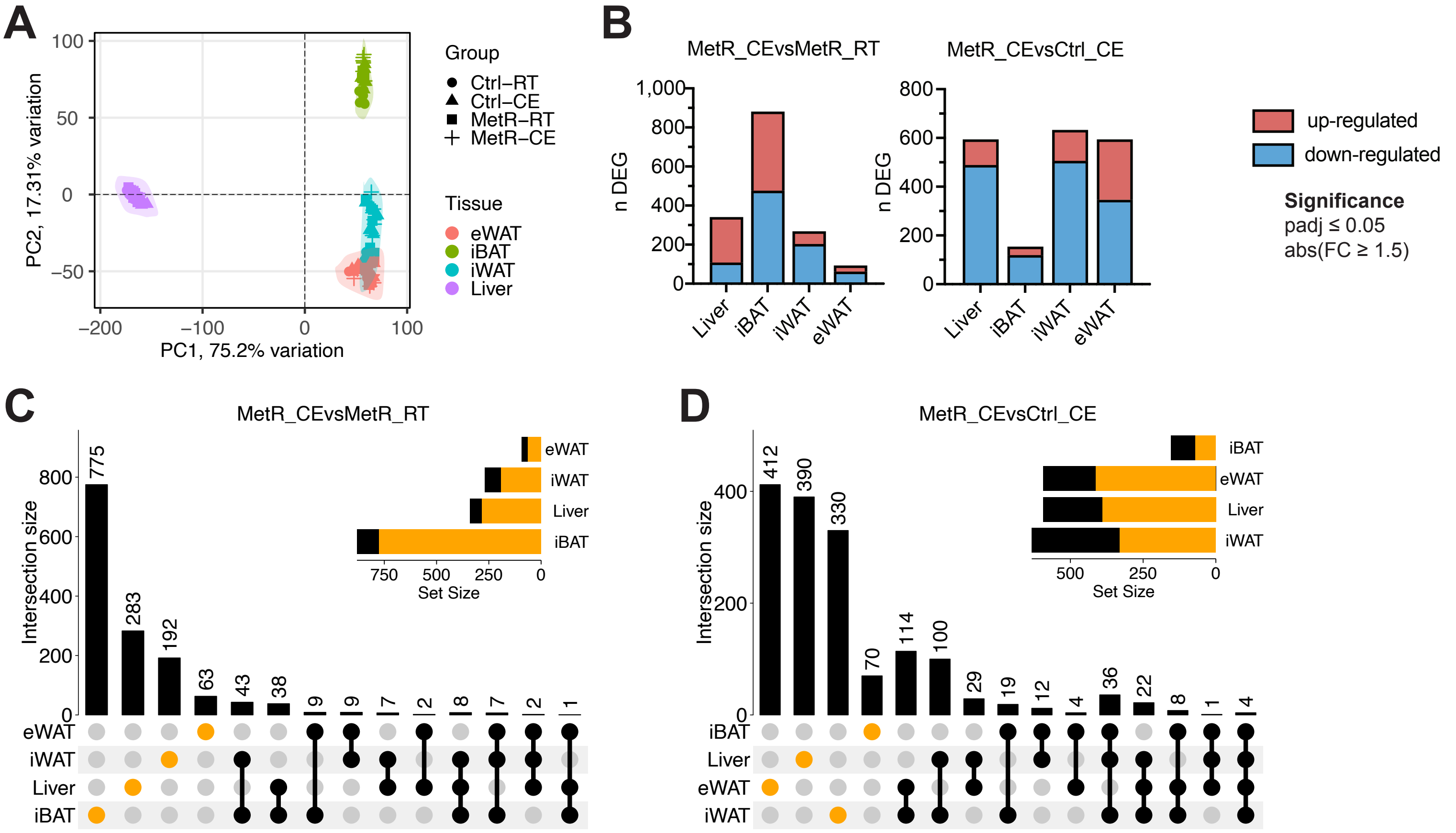

### Supplemental figure 3

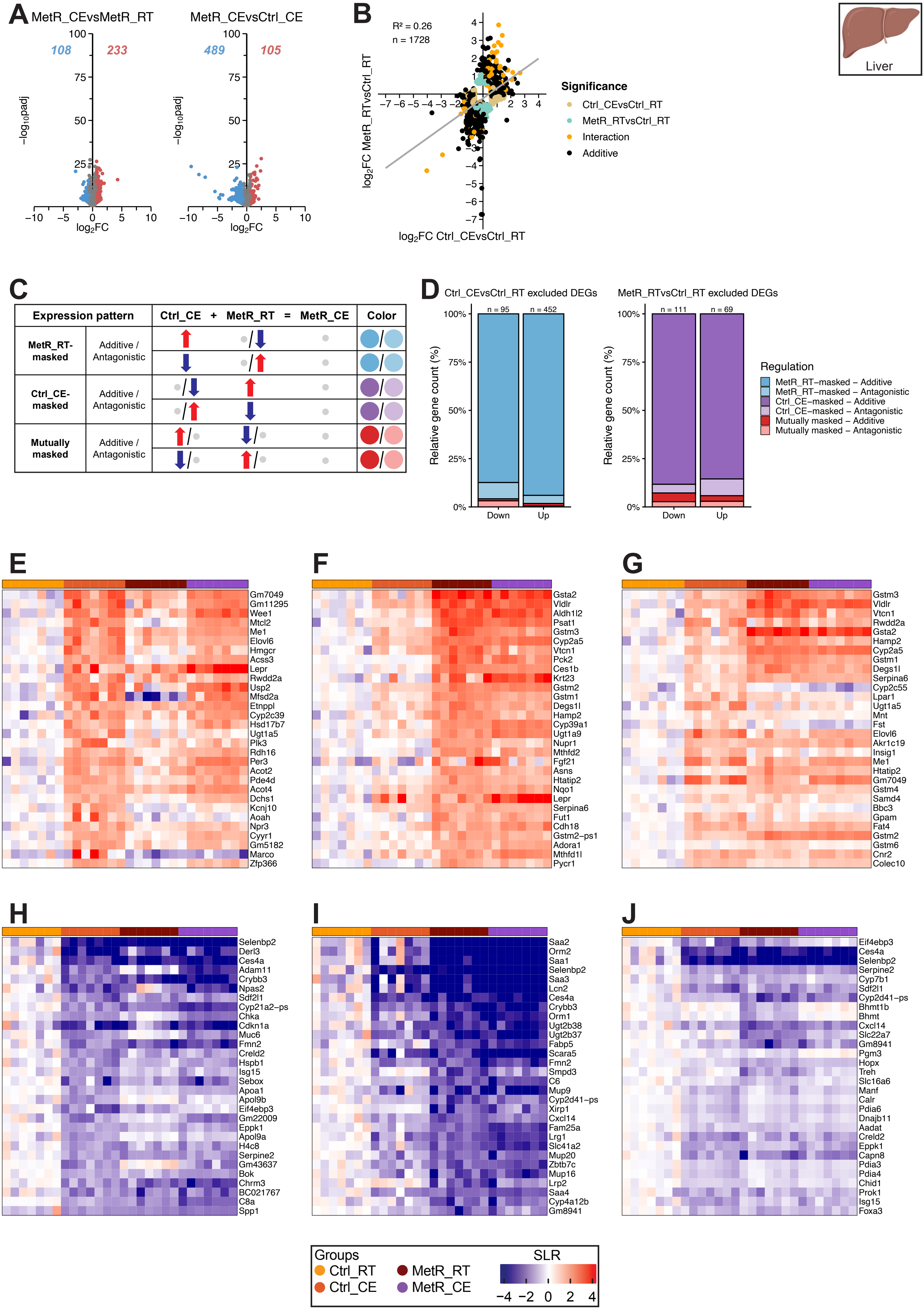

### Supplemental figure 4

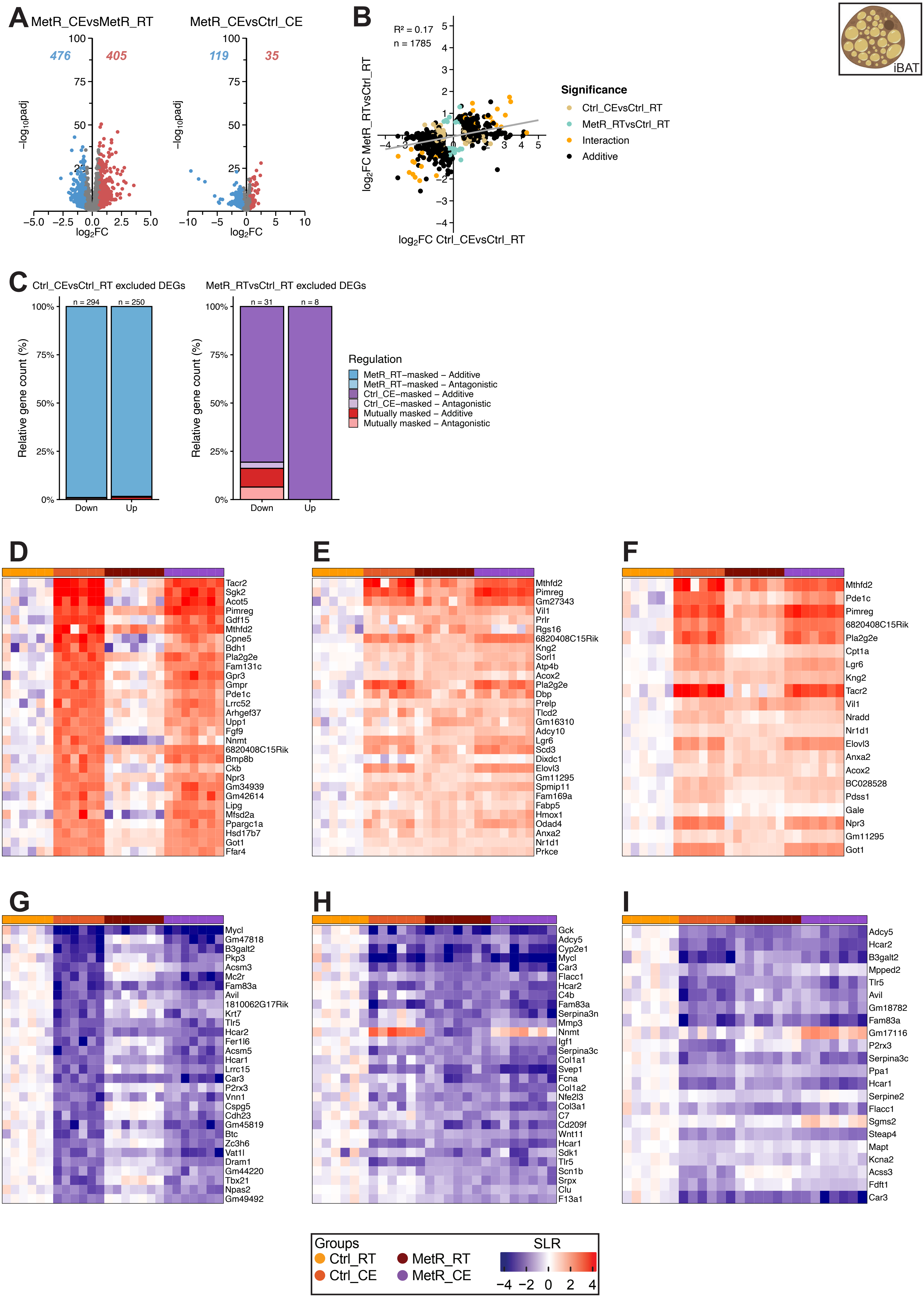

### Supplemental figure 5

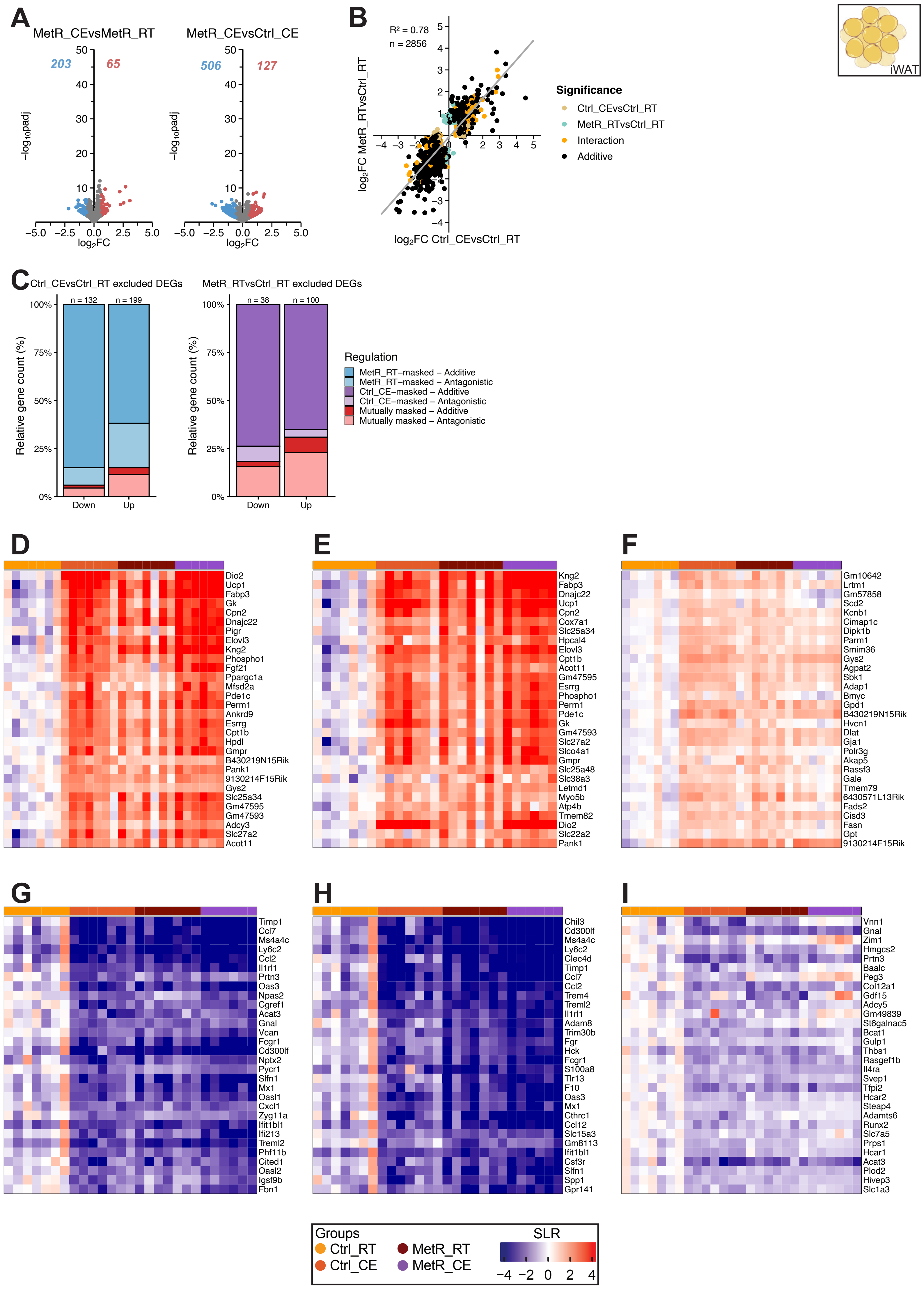

### Supplemental figure 6

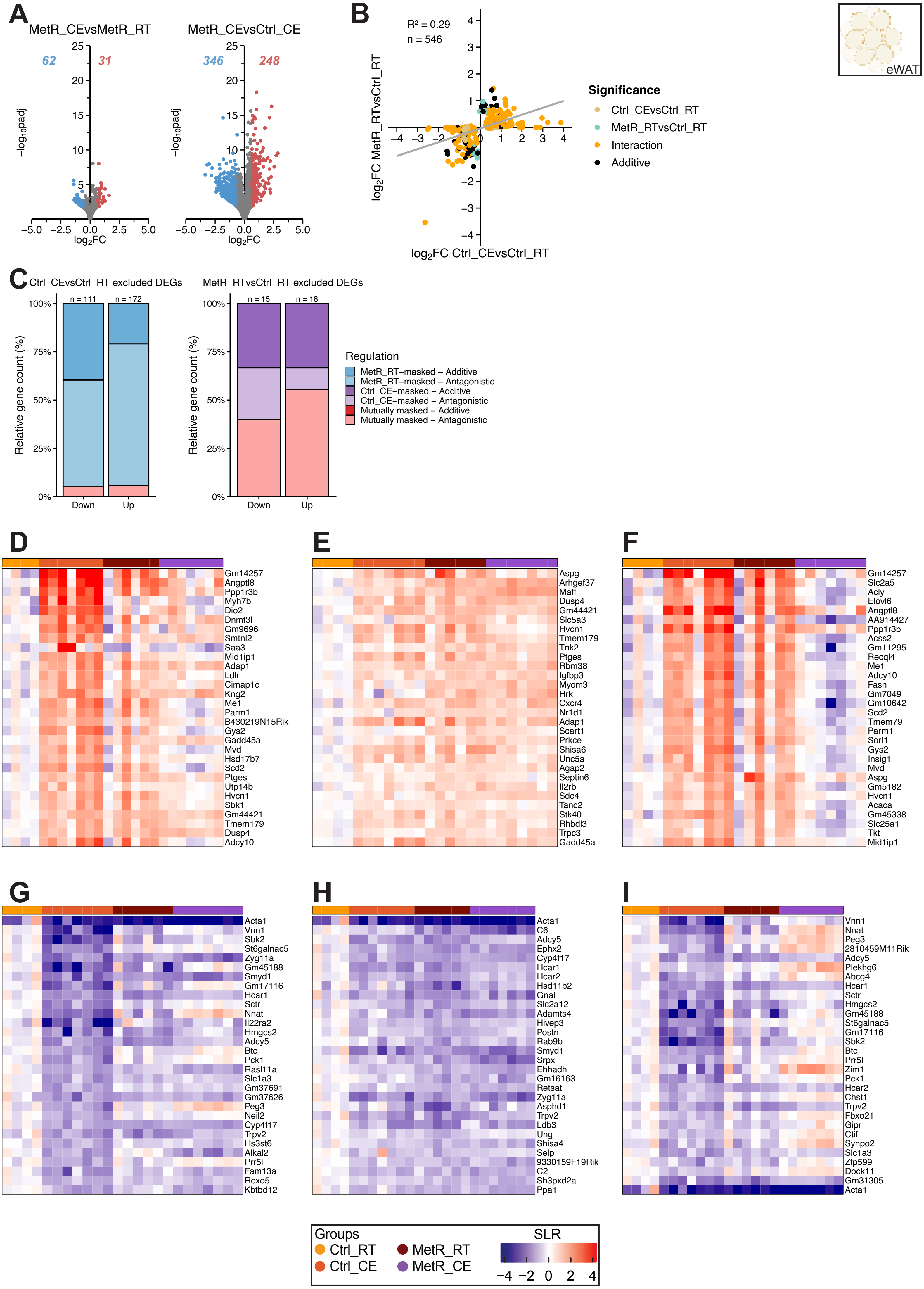
